# Diversification of ergot alkaloids and heritable fungal symbionts in morning glories

**DOI:** 10.1101/2021.06.09.447740

**Authors:** Wesley T. Beaulieu, Daniel G. Panaccione, Quynh N. Quach, Katy L. Smoot, Keith Clay

**Affiliations:** Department of Biology, Indiana University, Bloomington, IN, USA; Jaeb Center for Health Research, Tampa, FL, USA; Division of Plant & Soil Sciences, West Virginia University, Morgantown, WV, USA; Department of Ecology and Evolutionary Biology, Tulane University, New Orleans, LA, USA

## Abstract

Heritable microorganisms play critical roles in life cycles of many macro-organisms but their prevalence and functional roles are unknown for most plants. Bioactive ergot alkaloids produced by heritable *Periglandula* fungi occur in some morning glories (Convolvulaceae), similar to ergot alkaloids in grasses infected with related fungi. Ergot alkaloids have been of longstanding interest given their toxic effects, psychoactive properties, and medical applications. Here we show that ergot alkaloids are concentrated in four morning glory clades exhibiting differences in alkaloid profiles and are more prevalent in species with larger seeds than those with smaller seeds. Further, we found a phylogenetically-independent, positive correlation between seed mass and alkaloid concentrations in symbiotic species. Our findings suggest that heritable symbiosis has diversified among particular clades by vertical transmission through seeds combined with host speciation, and that ergot alkaloids are particularly beneficial to species with larger seeds. Our results are consistent with the defensive symbiosis hypothesis where bioactive ergot alkaloids from *Periglandula* symbionts protect seeds and seedlings from natural enemies, and provide a framework for exploring microbial chemistry in other plant-microbe interactions.

## Introduction

Microbial biochemistry plays a protective role in many macro-organisms where symbiotic hosts acquire higher fitness than non-hosts in the presence of natural enemies. This stands in contrast to nutritional symbioses where the microbial symbionts provide limiting resources to the host as in symbiotic N-fixation in legumes and provisioning of essential amino acids in aphids^1^. In many protective symbioses, the microbial symbionts are transmitted to offspring through the maternal lineage. For example, beewolf wasps (Hymenoptera) harbor antibiotic-producing symbiotic bacteria that protect larvae from fungal infections^2^, and intracellular bacteria producing kahalalide compounds protect their algal hosts against predation^3^. In many grasses (Poaceae), heritable fungal symbionts produce bioactive ergot alkaloids with anti-herbivory properties^4^. These defensive symbioses represent rich sources of natural products, and tractable systems for investigating the dynamics of interspecific interactions. More generally, understanding the origin, distribution and function of heritable symbioses provide insights into the evolution of major groups of eukaryotes.

Some morning glory species (family Convolvulaceae) contain high concentrations of ergot alkaloids (EAs) similar to those found in grasses infected with fungi in the family Clavicipitaceae^4,5^. EAs, named for ergot fungi (*Claviceps* spp.), are characterized by a common tetracyclic ergoline structure with a variety of complex side chains, and were first isolated from morning glory seeds in 1960^6^ based on their use as entheogens by indigenous peoples in Mexico^7,8^. EAs have been of longstanding interest given their toxic effects on humans and animals, medical applications and psychoactive properties^9^. For example, the EA lysergic acid provides the precursor for lysergic acid diethylamide (LSD). Much previous research has focused on increased herbivore and stress resistance of symbiotic grasses containing these fungal alkaloids^10^.

Morning glories are commonly considered as species of *Ipomoea* but this large genus is not monophyletic^11^, and a more accurate concept of morning glories is members of the tribe Ipomoeeae (morning glories with spiny pollen)^12^. Species are distributed across the world’s tropical and subtropical regions, are commonly found in disturbed habitats and thrive in open ecotonal areas such as forest and river edges^13^. Morning glories are important for humans as food crops (*I. batatas*, sweetpotato; *I. aquatica*, water spinach), agricultural weeds (*I. purpurea*, *I. triloba*) and ornamental flowers (*I. alba*, *I. tricolor*), and are an emerging model system for evolutionary studies^14^.

While it was known that *Claviceps* and related species produce EAs in their grass hosts, morning glories were initially thought to produce EAs endogenously until researchers^15^ treated *I. asarifolia* with fungicide and observed that EAs and characteristic epiphytic fungal colonies disappeared, suggesting that EAs in morning glories are also produced by symbiotic fungi. The EA biosynthetic gene, *dma*W, and nuclear rDNA sequences typical of Clavicipitaceae were identified in the fungus^16^, which infected plants systemically and was vertically transmitted through seeds^17,18^. Epiphytic fungal colonies can be observed in many symbiotic morning glory species on the adaxial surface of young leaves and closely associated with oil glands, providing a potential pathway for fungal nutrition and horizontal transmission, which has not been experimentally observed. This new fungal genus associated with Convolvulaceae was named *Periglandula* and genetic analysis grouped the fungi within the Clavicipitaceae^19^.

Prior to the discovery of *Periglandula* symbiosis, 42 species of morning glories out of 98 tested^5^ were reported to contain EAs. All reports were from the tribe Ipomoeeae, which includes the genera *Argyreia*, *Stictocardia*, *Turbina*, and the paraphyletic genus *Ipomoea*^20,21^. Two *Periglandula* species have been described to date based on their host associations and exhibit considerable genetic differentiation between them^19^. Efforts to cultivate *Periglandula* have been thus far unsuccessful, suggesting that some host products are necessary for fungal growth. Recent studies^22–24^ reported the co-occurrence of *Periglandula*, ergot alkaloid synthesis genes, and EAs in seeds of 10 additional morning glory species beyond the initial two described. Notably, *Periglandula* symbiosis with Convolvulaceae is macroscopically asymptomatic with hereditary (vertical) transmission through seeds and EA production at concentrations up to 1000X higher than in grasses infected with symbiotic *Epichloë* species. Moreover, morning glories are the only dicotyledonous plants known to contain EAs from fungal symbionts that have no known sexual reproduction, spore production or pathogenic relatives, unlike related Clavicipitaceae of grasses.

Given the vertical transmission of EAs and *Periglandula* through host seeds, we hypothesize that both symbiosis and EA chemistry will vary among host lineages through evolutionary diversification and speciation, and reflect the ecological costs and benefits of symbiosis. To address this hypothesis, we 1) quantified EAs in seeds of 210 morning glory species from diverse, worldwide herbarium collections in order to determine the global distribution and diversity of *Periglandula* symbiosis, 2) obtained phylogenetically-informative ITS sequences from GenBank and determined the distribution of EA-positive (EA+) and EA− species across the host phylogeny to elucidate evolutionary history of the symbiosis, and 3) evaluated whether EAs and *Periglandula* symbiosis vary with plant life-history traits to understand potential selective benefits of symbiosis for the host. Our results provide insights into the evolution and chemistry of heritable fungal symbiosis, and the diversification of EAs in dicotyledonous plants.

## Results

### Distribution of EAs in morning glory species

One-quarter of morning glory species that we evaluated (53 of 210 species with mature seeds from herbarium specimens; Fig. 1) contained EAs in one or more samples, including 36 previously unreported species (Table 1, Supplementary Fig. 1, Supplementary Data 1). Considering that we were unable to obtain seed samples from herbarium specimens of many morning glory species, the total number of EA+ species is certainly larger. *Ipomoea batatoides* had the highest total mean ergot alkaloid concentration in seeds with *I. hildebrandtii*, *I. kituiensis*, *I. racemosa*, *I. urbaniana*, and three of four *Stictocardia* species all having greater than 1,000 μg/g mean concentration (Supplementary Data 1). Cycloclavine, which was previously known only from one morning glory species (*I. hildebrandtii*), was detected in three species (*I. cicatricosa*, *I. hildebrandtii* and *I. killipiana*). The EAs chanoclavine, ergine, ergobalansine and ergonovine were widely distributed and detected in two-thirds or more of all EA+ species. We also found a mix of both positive and negative accessions for 13 of 19 EA+ species with three or more accessions, with six species consisting of only EA+ specimens (Supplementary Data 1). Species for which we had at least three accessions, and contained EAs in at least one sample, had alkaloids in 67% of the samples on average. Our results may underestimate the distribution and prevalence of EAs and *Periglandula* symbiosis given that a number of species were represented by only a single EA− sample (Supplementary Fig. 2a).

**Figure 1.**
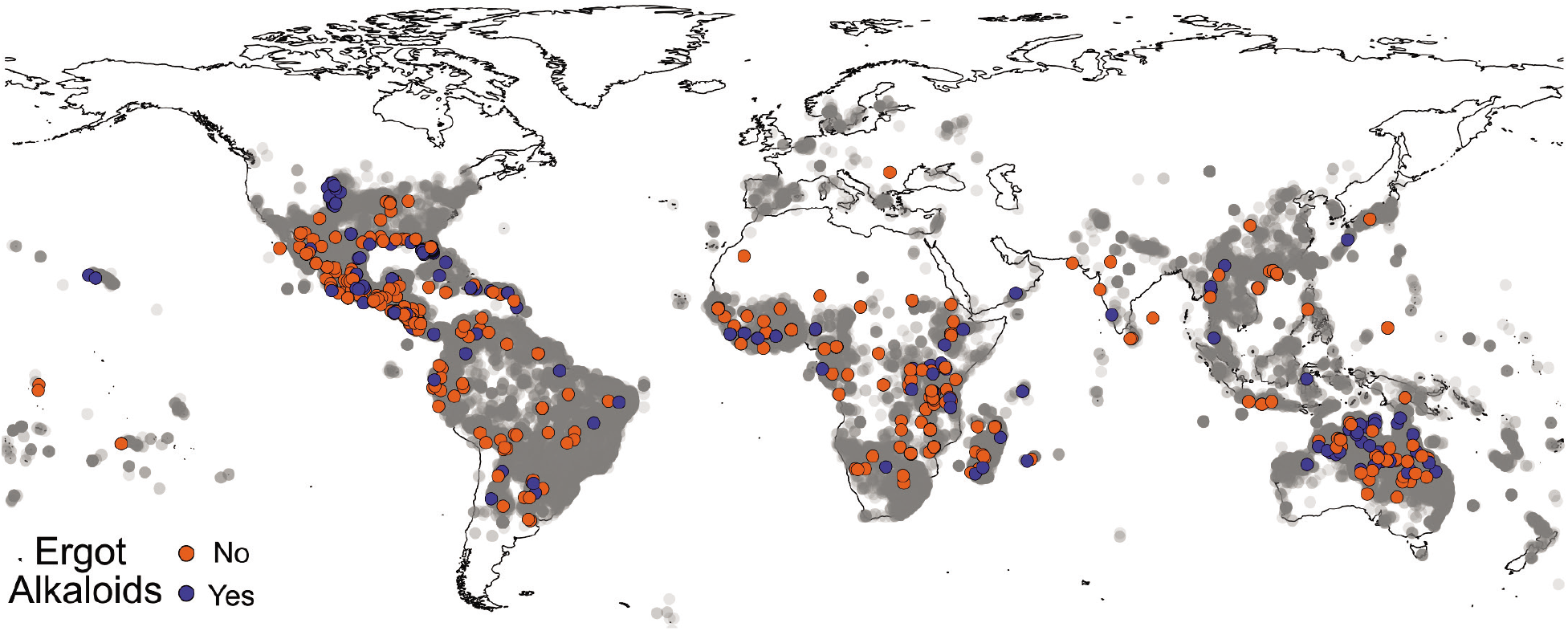
Geography of samples. Global distribution of samples tested for EAs (n=723). Grey circles represent herbarium specimens from *Argyreia*, *Ipomoea*, *Stictocardia*, and *Turbina* (now *Ipomoea*) species from the Global Biodiversity Information Facility^83–86^ to illustrate the geographical distribution of the tribe Ipomoeeae with higher concentrations of sample in darker areas.

**Table 1.** EA-positive species and average alkaloid composition. Species are listed by ITS phylogenetic order with double lines separating species from different origins of symbiosis (See Fig. 3). Values are raw ergot alkaloid concentrations (ug/g). The last six species were not included in our phylogeny.

| Species | chano-<br>clavine | cyclo-<br>clavine | agro-<br>clavine | lysergol | ergo-<br>novine | LAH<sup>±</sup> | ergine | ergosine | ergo-<br>balansine |
| --- | --- | --- | --- | --- | --- | --- | --- | --- | --- |
| *I. parasitica* | 97.6 | 0.0 | 0.0 | **565.4** | 10.6 | 0.0 | 0.0 | 106.3 | 139.2 |
| *I. leucotricha*<sup>†</sup> | 2.4 | 0.0 | 0.0 | 0.0 | 0.0 | 0.0 | **28.6** | 0.0 | 0.0 |
| *I. stans*<sup>†</sup> | **19.9** | 0.0 | 0.0 | 1.7 | 1.5 | 0.0 | 0.0 | 0.0 | 0.6 |
| *I. tricolor* | 2.5 | 0.0 | 0.0 | 0.0 | 1.9 | 65.2 | **95.0** | 0.0 | 0.0 |
| *I. barbatisepala*<sup>†</sup> | 8.7 | 0.0 | 0.0 | 0.0 | 9.3 | 3.6 | **34.6** | 0.0 | 0.0 |
| *I. aculeata*<sup>†</sup> | 2.7 | 0.0 | 0.0 | 0.0 | 258.5 | 0.0 | **268.1** | 0.0 | 0.0 |
| *I. indivisa*<sup>†</sup> | 6.3 | 0.0 | 0.0 | 0.0 | 0.6 | 0.0 | 0.9 | 0.0 | **38.0** |
| *I. graminea*<sup>†</sup> | **1.2** | 0.0 | 0.0 | 0.0 | 0.0 | 0.0 | 0.0 | 0.0 | 0.0 |
| *I. aquatica*<sup>†</sup> | 7.4 | 0.0 | 0.0 | 0.0 | 5.6 | 15.1 | 15.6 | 0.0 | **56.1** |
| *I. prismatosyphon*<sup>†</sup> | 0.0 | 0.0 | 0.0 | 0.0 | 0.0 | 0.0 | 0.0 | 0.0 | **1.7** |
| *I. abrupta*<sup>†</sup> | 2.9 | 0.0 | 0.0 | 0.0 | 1.9 | 3.3 | 3.5 | 0.0 | **15.1** |
| *I. mauritiana*<sup>†</sup> | 0.0 | 0.0 | 0.0 | 0.0 | 2.3 | 0.0 | **2.7** | 0.0 | 2.2 |
| *I. batatoides*<sup>†</sup> | 172.1 | 0.0 | 0.0 | **2805.4** | 0.0 | 0.0 | 0.0 | 68.5 | 588.1 |
| *I. aurantiaca*<sup>†</sup> | 0.0 | 0.0 | 0.0 | 0.0 | 0.0 | 0.0 | 0.0 | 0.0 | **6.2** |
| *I. brassii*<sup>†</sup> | 2.9 | 0.0 | 0.0 | 0.0 | 0.1 | 5.6 | **6.8** | 0.0 | 4.5 |
| *I. gracilis* | 32.2 | 0.0 | 0.0 | 0.0 | 18.4 | 21.8 | 10.3 | 1.0 | **47.9** |
| *I. muelleri* | 5.6 | 0.0 | 0.0 | 0.0 | 4.4 | 23.3 | 13.6 | 0.0 | **50.2** |
| *I. argillicola* | 10.8 | 0.0 | 0.0 | 0.0 | **41.3** | 19.4 | 10.4 | 0.0 | 31.9 |
| *I. fimbriosepala*<sup>†</sup> | 10.0 | 0.0 | 0.0 | 0.0 | 4.0 | 14.2 | 5.1 | 0.0 | **69.6** |
| *I. setifera* | 3.4 | 0.0 | 0.0 | 0.0 | 2.6 | 16.4 | 15.3 | 0.0 | **30.1** |
| *I. pes-caprae* | 1.7 | 0.0 | 0.0 | 0.0 | 1.5 | 2.3 | **3.5** | 0.0 | 1.9 |
| *I. syringifolia*<sup>†</sup> | 2.6 | 0.0 | 0.0 | 0.0 | 0.6 | 0.0 | 1.5 | 0.7 | **137.1** |
| *I. amnicola* | 79.1 | 0.0 | 0.1 | 0.7 | 1.0 | 0.0 | 7.8 | 2.2 | **339.3** |
| *I. leptophylla* | **11.2** | 0.0 | 0.0 | 0.0 | 6.1 | 3.6 | 6.5 | 0.0 | 0.0 |
| *I. philomega* | 0.0 | 0.0 | 0.0 | 0.0 | 0.0 | 43.2 | **73.6** | 18.4 | 1.1 |
| *I. maurandioides*<sup>†</sup> | 50.1 | 0.0 | 0.0 | 0.0 | 27.1 | 26.9 | 16.2 | 0.0 | **201.1** |
| *I. incarnata*<sup>†</sup> | 5.6 | 0.0 | 0.0 | 0.0 | 5.5 | **7.4** | 4.4 | 0.0 | 0.0 |
| *A. osyrensis*<sup>†</sup> | 0.0 | 0.0 | 0.0 | 0.0 | 0.0 | 0.0 | 0.0 | **107.1** | 10.0 |
| *A. splendens* | 0.0 | 0.0 | 0.2 | 0.0 | 0.0 | 0.0 | 284.6 | **582.1** | 10.7 |
| *A. obtusifolia* | 16.5 | 0.0 | 0.0 | 0.0 | 0.6 | 0.0 | 7.5 | **67.1** | 2.7 |
| *A. henryi*<sup>†</sup> | 0.0 | 0.0 | 5.9 | 0.0 | 4.2 | 0.0 | **56.8** | 0.0 | 0.2 |
| *A. roseopurpurea*<sup>†</sup> | 8.0 | 0.0 | 0.0 | 0.0 | 0.0 | 0.0 | 25.7 | **143.8** | 4.3 |
| *I. porphyrea*<sup>†</sup> | 0.0 | 0.0 | 0.0 | 0.0 | 0.2 | 1.1 | 12.4 | **27.2** | 2.6 |
| *A. nervosa* | 0.5 | 0.0 | 0.0 | **18.9** | 14.0 | 0.0 | 10.1 | 2.6 | 0.0 |
| *I. sericosepala*<sup>†</sup> | 5.5 | 0.0 | 0.0 | **31.0** | 2.3 | 0.6 | 9.7 | 3.4 | 2.4 |
| *I. abutiloides* | 0.4 | 0.0 | 0.0 | 0.0 | 0.0 | 0.0 | 0.0 | **245.4** | 3.0 |
| *I. spathulata*<sup>†</sup> | 0.0 | 0.0 | 0.0 | 0.0 | 0.0 | 0.0 | 0.0 | **62.2** | 5.0 |
| *I. kituiensis*<sup>†</sup> | 15.4 | 0.0 | 139.1 | 88.4 | 3.7 | 13.2 | 23.3 | **910.1** | 3.5 |
| *I. cicatricosa*<sup>†</sup> | 9.9 | **211.2** | 0.0 | 0.0 | 1.1 | 14.1 | 5.6 | 0.0 | 0.3 |
| *I. urbaniana*<sup>†</sup> | 32.4 | 0.0 | 1.2 | 1.9 | 1.3 | 12.2 | 13.4 | **2773.0** | 23.0 |
| *I. adenioides*<sup>†</sup> | 0.0 | 0.0 | 0.0 | 0.0 | 0.0 | 0.2 | 0.8 | **6.1** | 1.9 |
| *S. beraviensis* | 2.6 | 0.0 | 0.0 | **1066.5** | 64.2 | 0.0 | 0.0 | 0.0 | 0.0 |
| *S. macalusoi*<sup>†</sup> | 2.6 | 0.0 | 0.0 | **1596.2** | 2.6 | 0.0 | 0.0 | 0.0 | 0.0 |
| *S. tiliifolia* | 34.7 | 0.0 | 0.0 | **1127.8** | 51.4 | 0.0 | 0.0 | 0.0 | 0.0 |
| *I. stenobasis*<sup>†</sup> | **5.2** | 0.0 | 0.0 | 0.0 | 0.4 | 0.0 | 2.4 | 0.0 | 0.0 |
| *I. pileata*<sup>†</sup> | 0.0 | 0.0 | 0.0 | 1.1 | 0.0 | 0.0 | 0.0 | **10.0** | 0.8 |
| *I. lindenii*<sup>†</sup> | 10.2 | 0.0 | 0.0 | 0.0 | 0.0 | 0.0 | 0.0 | 0.0 | **76.9** |
| *I. antonschmidii*<sup>†</sup> | 1.3 | 0.0 | 0.0 | 0.0 | 3.2 | 3.0 | 7.1 | 0.0 | **8.3** |
| *I. chrysochaetia*<sup>†</sup> | 0.0 | 0.0 | 0.0 | 0.0 | 0.0 | 0.0 | 0.0 | **19.5** | 0.0 |
| *I. hildebrandtii* | 3.7 | **1486.2** | 0.0 | 0.0 | 4.2 | 5.8 | 1.1 | 0.0 | 0.0 |
| *I. killipiana*<sup>†</sup> | 0.0 | **777.2** | 0.0 | 0.0 | 0.0 | 0.0 | 0.8 | 0.0 | 0.0 |
| *I. racemosa*<sup>†</sup> | 3.9 | 0.0 | 0.0 | 0.0 | **621.8** | 0.0 | 3.6 | 422.6 | 16.5 |
| *P. shirensis*<sup>†</sup> | 0.0 | 0.0 | 0.0 | 0.0 | 0.0 | 0.0 | 0.0 | **53.6** | 2.3 |

± lysergic acid α-hydroxyethylamide
† Previously unidentified species reported with ergot alkaloids
Bolded values notes the ergot alkaloid with the highest concentration in the species.

In addition to seeds obtained from herbarium specimens, we sampled mature seeds from natural populations of two widespread and easily accessible EA+ species to investigate heterogeneity of EAs among populations. These collections revealed that 21 of 21 populations of *I. leptophylla* from the northern Great Plains region of the United States and 38 of 38 populations of *I. pes-caprae* from Florida beaches contained EAs, indicating that *Periglandula* symbiosis is ubiquitous in these regions. EA concentrations were significantly lower in *I. pes-caprae* (mean 12.6 ug/g) than in *I. leptophylla* (mean 28.2 ug/g; (*t*(115.4) = −8.62, *p*<0.001). Unexpectedly, EA concentrations in *I. pes-caprae* were 43% higher in Gulf Coast populations compared to Atlantic Coast populations (*t*(36.0) = −3.91, *p*<0.001), potentially reflecting higher nutrients in the Gulf of Mexico or genetic differentiation between the east and west sides of the Florida peninsula.

### Latitude, sample age, and EAs

We tested whether latitude of origin and age of herbarium specimens affected the likelihood of detecting EAs. There was no significant effect of latitude on the presence of EAs (*p*=0.46) or the total concentration of EAs among EA+ species (*R^2^*=0.004, *p*=0.65). Herbarium specimens ranged between five and 179 years old, with the majority between 20 and 60 years old (Supplementary Fig. 2b), but we found that EA concentrations were not correlated with sample age (*R^2^*=0.005, *p*=0.43, Supplementary Fig. 3), indicating that EAs persist in seeds over extended periods.

### Comparison with previous survey

Eich^5^ previously categorized 98 morning glory species as “unambiguously positive”, “apparently devoid”, or having “conflicting reports” for EAs, based primarily on publications by other groups using variable methodologies and plant tissues. Our results from 68 of those 98 species agreed with the Eich classifications (Supplementary Data 1), but the majority of species represented in our seed samples were not included in Eich^5^. We identified four EA+ species listed as “apparently devoid” (*I. adenioides*, *I. batatoides*, *I. graminea* and *I. mauritiana*), although only one accession of each species contained EAs (N=1, 3, 2 and 6 accessions respectively). There were also eight *Ipomoea* species listed as EA+ that we found to be EA-, and we detected no alkaloids in 13 of 14 species listed as “contradictory”^5^. In the one other contradictory species, *I. aquatica*, we found that two of four accessions contained EAs. Polymorphism within species may contribute to some of these discrepancies, as may methodological differences among the numerous studies cited by Eich^5^ and potential misidentifications of species.

### EAs distribution in host phylogeny

To evaluate the occurrence of EAs in relation to host phylogeny, we obtained published ITS sequences for 183 of our 210 sampled species, plus four outgroup species and 19 additional species reported by Eich^5^ (Supplementary Table 1, Supplementary Data 2). We found significant phylogenetic signal for the presence of EAs (Pagel’s *λ*= 0.8, *D*=0.059, *p*<0.001), suggesting that species with EAs are more closely related than expected by chance (Figs. 2, 3). By combining morning glory systematics^25,26^ with recent molecular results^21,27^, we found that most EA+ species occurred in four major clades, along with other single EA+ species occurrences dispersed through the phylogeny (Figs. 2, 3, Supplementary Figs. 4, 5). The largest clade of 22 species sampled here consists of three lineages: *Argyreia*, two ‘*Turbina*’ species, and the African clade Poliothamnus (ATP clade, Fig. 3; note that recent taxonomic treatments have dissolved the genus *Turbina* into *Ipomoea*^28^). It is well-represented in Asia but also includes African and South American distributions. The Pes-caprae clade (18 species) is largely New World but also includes Australian species and pantropical *I. asarifolia*, *I. fimbriosepala*, and *I. pes-caprae*. The other clades are *Stictocardia* (four species, largely African) and Tricolores (four species, the Americas). Phylogenetic analysis also suggests that EAs are absent from several large clades (Figs. 2, 3). For example, our samples from the Batatas clade, which includes sweetpotato, and the Jalapae clade, are entirely EA− (Fig. 3, Supplementary Fig. 4). A distinct heritable fungal symbiont that produces the toxic indolizidine alkaloid swainsonine occurs in *I. carnea*, *I. costata* and other species in the Jalapae and related clades^29,30^, potentially reflecting selection against redundant functions of multiple symbionts or their increased physiological costs such that only one endosymbiont can persist within a single host species^31,32^. In other clades, most species are EA− with occasional EA+ taxa (e.g., *I. batatoides*, *I. mauritiana*). Conversely, sporadic EA− species occur within the largely EA+ Pes-caprae and ATP clades (e.g., *I. pandurata*, *A. capitiformis*; Fig. 3). These instances suggest that both occasional gains and losses of symbiosis have occurred within clades. Overall, our results suggest that the Ipomoeeae common ancestor was not symbiotic, but rather that symbiosis arose independently in one or a few clades and diversified via vertical transmission and host speciation, with occasional host jumps through horizontal transmission by a yet unidentified mechanism, potentially through the transfer of epiphytic colonies (Supplementary Fig. 6).

**Figure 2.**
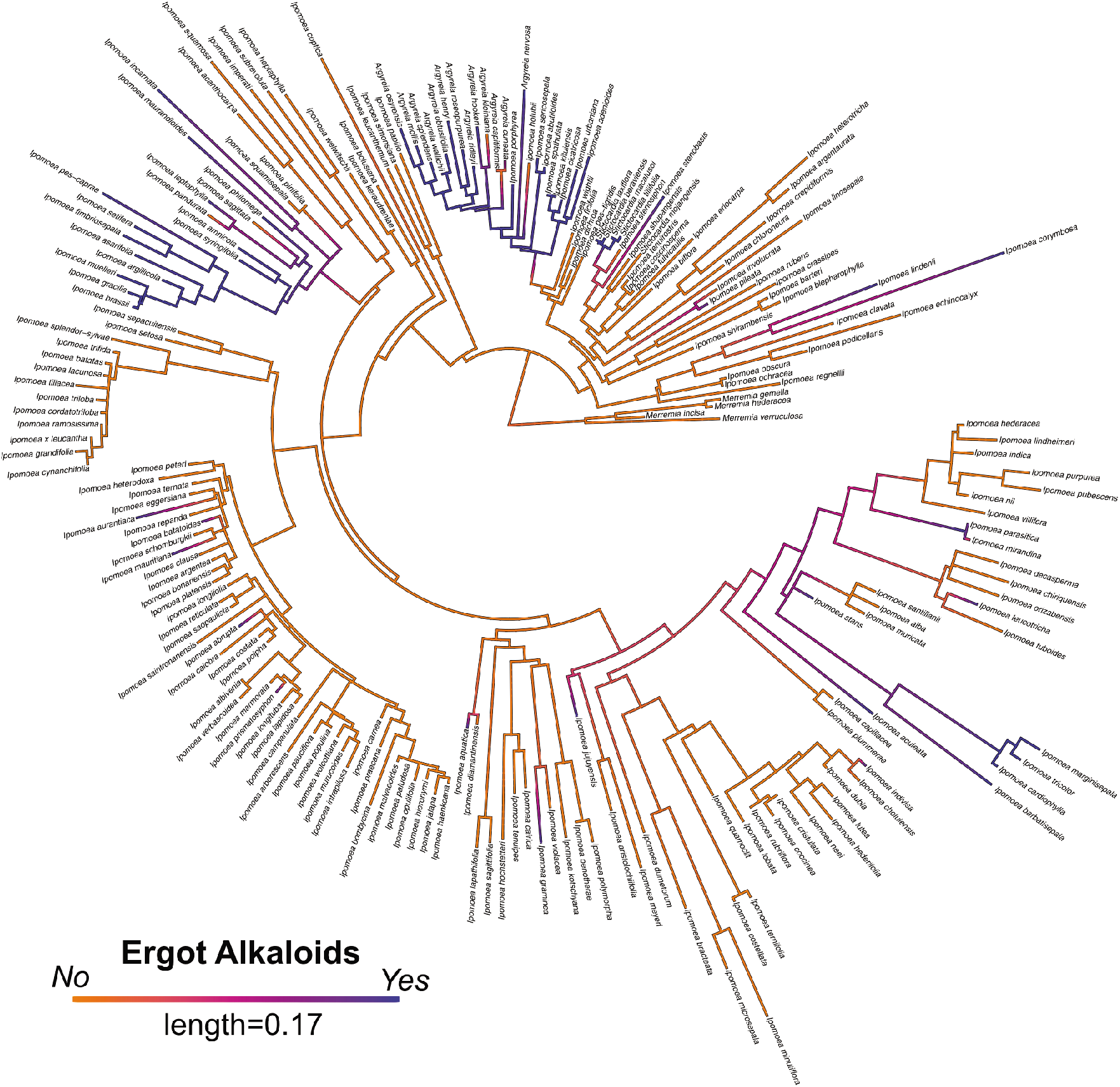
ITS phylogeny of morning glories. Maximum likelihood ITS phylogeny of morning glories (n=206) with ancestral state reconstruction of EA presence by density map based on 1000 stochastic character maps. Branch color represents the probability of the character state. Length of the legend equals units of substitution per site.

**Figure 3.**
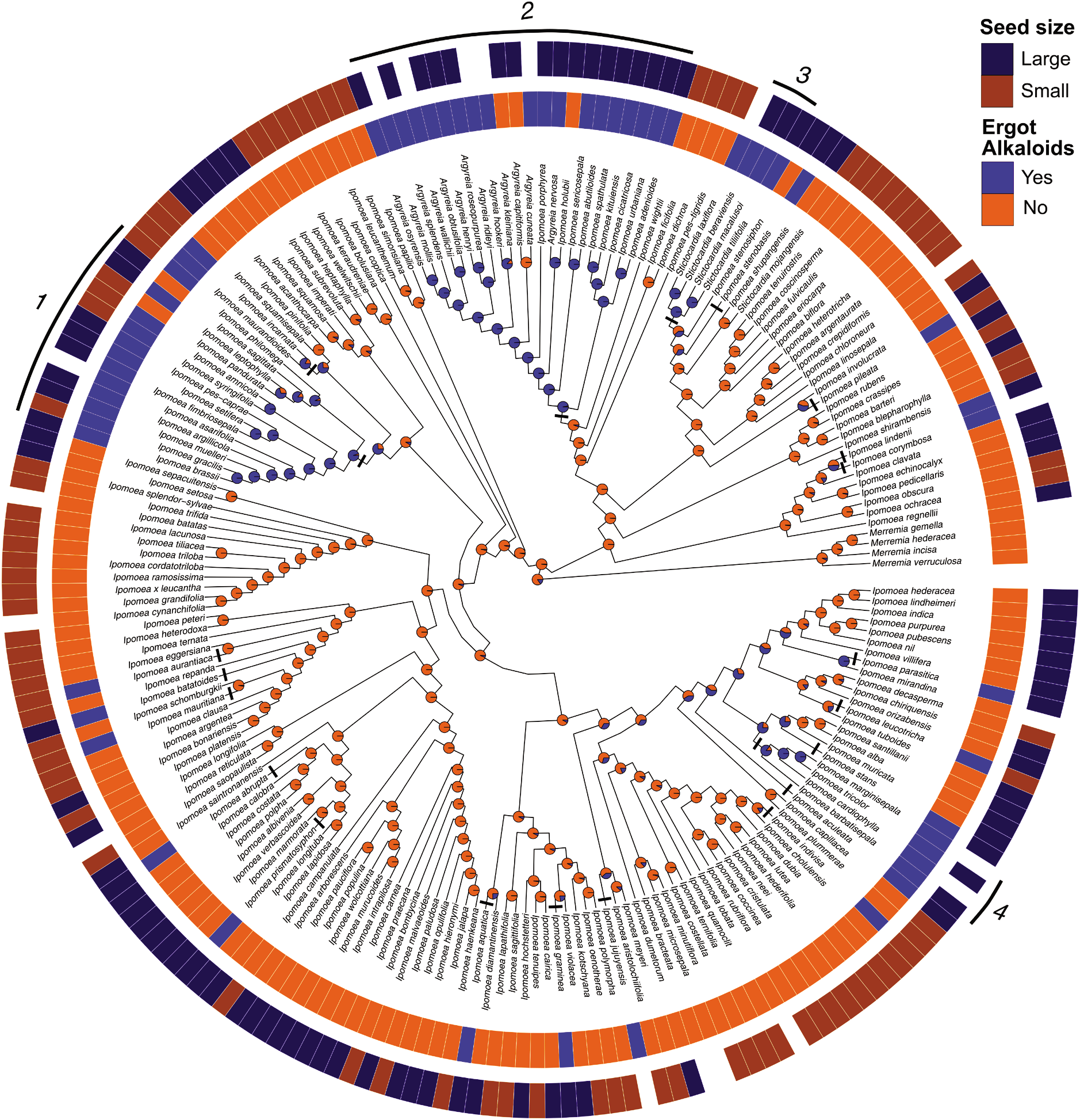
Ancestral state reconstruction of EA presence. Phylogenetic distribution of EA presence and seed mass. Species lacking seed mass data are represented by blank spaces. Pie charts at nodes indicate the relative likelihoods of the alkaloid ancestral character state. Black bars represent the origins of symbiosis. Lines and numbers indicate major clades of EA+ species: 1) Pes-caprae, 2) ATP, 3) Stictocardia, 4) Tricolores.

### EAs chemistry and host phylogeny

For heritable symbionts lacking contagious spread, related host species should support related fungal symbionts with similar EA profiles through descent from common ancestors. Research with *Epichloë*-infected grasses indicates that the identities of EAs are determined by the fungus and not the host plant^33^. We detected nine EAs that occurred in 59 combinations (Table 1), and conducted phylogenetic principal component analysis (PCA) on the average concentrations of different alkaloids per species to determine whether EA profiles map onto host phylogeny. Analysis of similarity indicated significant dissimilarity of EA profiles among major clades (*R*=0.31, *p*=0.001; Fig. 4a, Supplementary Fig. 7a). For example, EAs in species from the Pes-caprae clade were dominated by ergobalansine while species in the ATP clade contained primarily ergosine (Table 1). Alkaloid profiles also differed significantly among species within the Pes-caprae clade (*R*=0.52, *p*=0.001, Fig. 4b) and within the ATP clade (*R*=0.53, *p*=0.001, Supplementary Fig. 7b).

**Figure 4.**
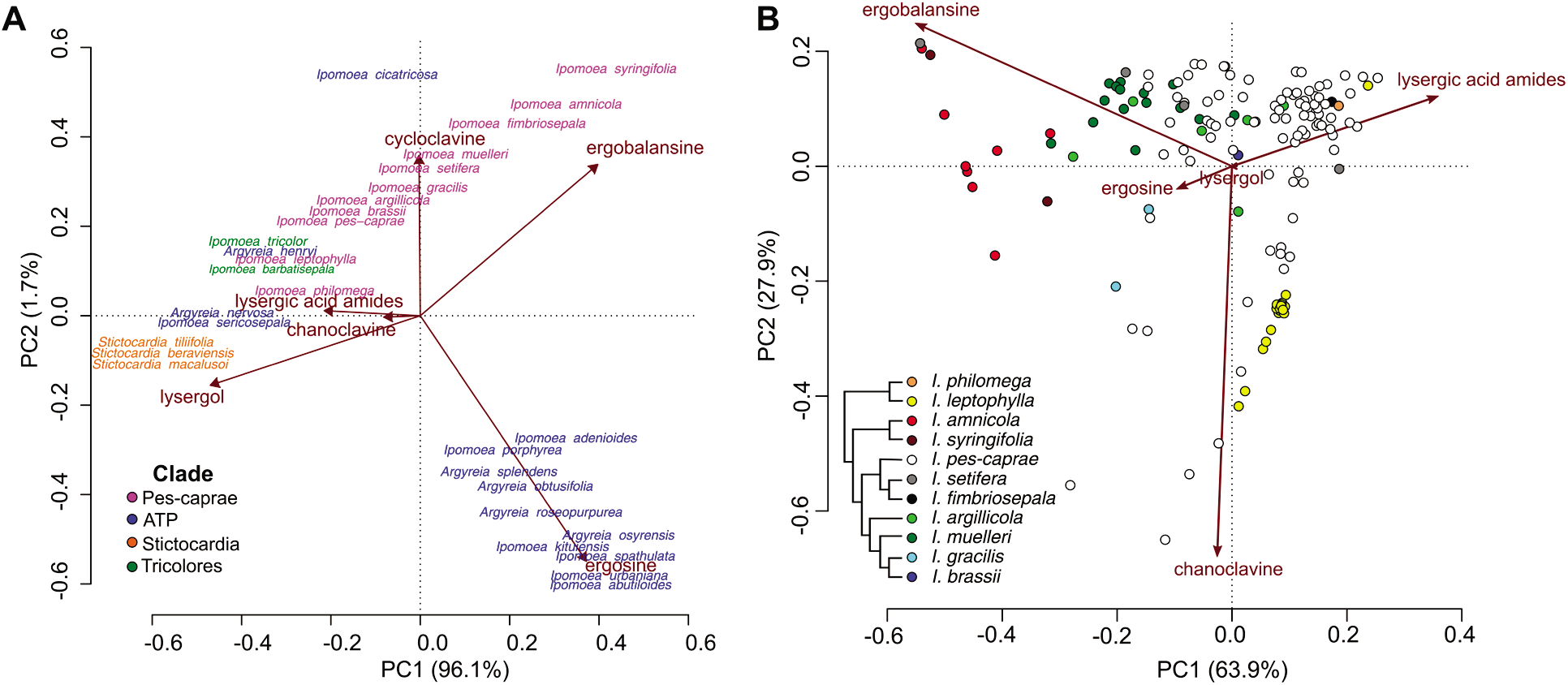
Clustering of alkaloid profiles in EA+ species. **(a)** Phylogenetic PCA of EA+ species colored by the four major clades. EA values are averaged for each species. **(b)** PCA of EA+ species in the Pes-caprae clade.

### Seed mass and EAs

We also measured seed mass from herbarium samples as a proxy for plant life history where larger-seeded species have lower growth rates and are longer lived than smaller-seeded species^34^. Seeds of EA+ species were significantly larger than seeds of EA− species (50.7 mg vs. 37.5 mg, *t*(118.1)=-3.98, *p*<0.001, Fig. 5a), where seed mass varied 100-fold among species (Supplementary Data 1). To further explore this relationship, we transformed seed mass to a binary state and found a significant excess of large-seeded species containing EAs (*X^2^*=13.3, *p*<0.001, Fig. 5b). We then performed Pagel’s test^35^, which revealed a significant correlation between seed mass and EAs after accounting for host phylogeny (*p*<0.05, Fig. 3, Supplementary Fig. 8). Independent of EAs, there was a significant phylogenetic signal in seed mass (*λ*= 0.79, *p*<0.001), suggesting that closely related species have more similar seed mass. Finally, we conducted phylogenetic regression^36^, which demonstrated that EA concentrations in EA+ species increased significantly with seed mass after accounting for phylogenetic correlation (*p*=0.01, Fig. 5c). Thus, EA+ species have larger seeds than EA− species and EA concentrations increase with increasing seed mass in EA+ species.

**Figure 5.**
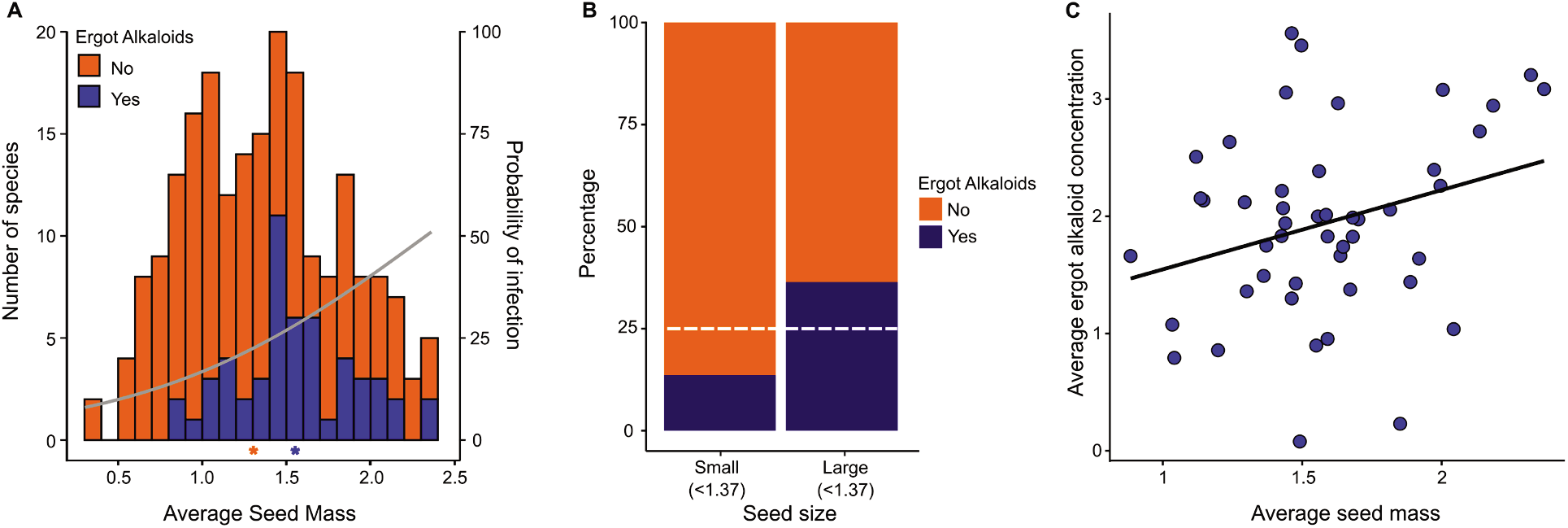
Relationship of seed mass and EA concentrations. Seed mass and EA concentrations are log10-transformed. **(a)** Number of species with (n=53) or without (n=157) EAs by mean seed mass. Stars along the x-axis indicate the mean seed mass of each group. Binomial regression line shows the probability of infection based on mean seed mass. **(b)** Proportion of small and large-seeded species with or without EAs. Dashed lines represent the expected number of EA+ species if the proportion of EA+ species is equivalent across seed mass. **(c)** Phylogenetic regression of seed mass and EA concentration in EA+ species.

## Discussion

The potential origins of *Periglandula* symbiosis include horizontal transfer from Clavicipitaceae-infected grasses or from clavicipitaceous entomopathogens of hemipteran parasites of morning glories^37,38^. Notably, clavicipitaceous entomopathogens also exhibit diverse bioactive chemistry^38^. Phylogenetic studies clearly indicate that *Periglandula* is distinct from grass- and insect-infecting Clavicipitaceae^24,39^, supporting its ancient origins. Based on previous research^21,40^, we estimate the *Periglandula* symbiosis arose in the ATP clade c. 15 MYA. By comparison, *Claviceps*-infected grasses have been reported from Cretaceous amber 100 MYA^41^. Once established, there must have been occasional host shifts among clades of morning glories but wholesale horizontal spread is not supported by the phylogenetic patterns reported here. Phylogenetic patterns of heritable symbiosis also occur within other plant families. For example, fungal symbiosis and swainsonine production in locoweeds (Leguminosae) is limited to three host genera (out of 750), and *Epichloë* endophytes occur only in cool-season grasses in the subfamily Pooideae^4^.

What are the costs and benefits of EAs and *Periglandula* symbiosis? Heritable symbionts transmitted through seeds should be mutualistic given that they will not persist if they reduce host fitness^42^. While we do not have direct evidence that seed survival, germination, or resistance to predation are enhanced by EAs in morning glories, two experimental studies of *Epichloë*-infected grasses where EAs occur in seeds reported that seed-harvesting ants exhibit a significant preference for symbiont-free seeds over symbiont-infected seeds^43,44^. Similarly, multiple passerine bird species referred symbiont-free grass seeds over symbiont-infected seeds^45^. Further, increased resistance to herbivory has been demonstrated in cool-season grasses symbiotic with *Epichloë* compared to symbiont-free conspecifics^46^, and pure EAs, including many found in morning glories, applied to food plants reduced insect growth^47^. In morning glories, EAs are concentrated in seeds where they could play a role against seed and seedling predators. For example, many morning glories are attacked by bruchine beetles^48^ (Chrysomelidae) where larvae feed within developing seeds. Poisoning of livestock following grazing on EA+ morning glory species has also been reported^49^. In *I. tricolor*, EAs in seeds are redistributed to host roots following germination^50^, and young EA+ plants are significantly more resistant to root-knot nematodes than EA− plants^32^. However, EA+ plants had reduced growth compared to EA− plants in the absence of nematodes, suggesting a physiological cost of symbiosis for host plants. Our results imply that EAs are disproportionately beneficial to species with larger seeds, which represent a larger investment by the maternal plant. This benefit may be direct where larger seeds and seedlings are subject to greater predator and pathogen pressure, or indirect where large seed mass is correlated with other plant life history traits like slower growth rate and longer life-span. More broadly, our results confirm the strong linkage between heritable fungal symbiosis and bioactive toxins across multiple plant families^51^, and suggest host-symbiont coevolution manifested as larger seeds with higher EA concentrations.

One limitation of this study is that we could only test 210 of 820 species in the Ipomoeeae because most herbarium samples did not contain mature seeds, given that specimens are typically collected during the diagnostic flowering stage. Extrapolation of our results to all morning glories suggests that there could be 150 or more additional EA+ host species yet to be detected. This may be a conservative estimate given that EA− species represented by one sample here could prove EA+ with additional samples (Supplementary Data 1, Supplementary Fig. 2a). Integrating our EA data into the larger dataset of *Ipomoea* species presented by Munoz et al^.27^ potentially identifies other likely EA+ species (Supplementary Data 3). For example, the Pes-caprae clade may contain other EA+ members such as *I. procurrens* and *I. paludicola* based on Munoz’s phylogeny. Independent of sampling depth, the ITS phylogeny could be expanded to a multigene phylogeny in the future with additional sequencing. At present, most of the available DNA sequence data for morning glories come from a single large published ITS phylogeny with over 400 species^27^. Another possible limitation arises if there are EA− *Periglandula* symbionts, which would underestimate symbiosis based on EA detection, although there is no evidence of this. For example, a previous study^24^ found that EAs and *Periglandula* sequences co-occurred in all eight morning glory species examined. Culturing of *Periglandula* has not yet been achieved, but would allow more detailed investigations of the symbiont, EA production, and host specificity.

Our results demonstrate that morning glories containing ergot alkaloids produced by *Periglandula* fungal symbionts are distributed worldwide and we report 36 previously unreported species containing EAs, suggesting that the numbers of host taxa engaged in *Periglandula* symbiosis will grow with additional research. Further, the EA+ species are concentrated in particular clades, consistent with the hypothesis that the *Periglandula* symbionts are vertically transmitted within clades with infrequent host jumps to distantly-related species. EA+ clades exhibited significant differences in EA profiles, reflecting genetic differences among *Periglandula* in EA biosynthetic pathways. EA+ species also had larger seeds with significantly higher EA concentrations, supporting the hypothesis that the symbiont provisions EAs to critical life-history stages for both the host and fungus. An important future direction will be to evaluate the importance of EAs and *Periglandula* symbiosis on host fitness in experimental or natural populations. Understanding why bioactive EAs and *Periglandula* symbiosis have diversified in certain clades and their relationship to plant life-history traits will provide a predictive framework that can be applied more broadly to other heritable fungal symbioses in plants.

## Methods

### Taxon Sampling

We sampled mature seeds from a total of 210 morning glory species from 723 herbarium sheets, where species identity, mean alkaloid content, mean seed mass, and mean collection latitude are reported by species for each herbarium specimen (Supplementary Data 1, Supplementary Data 4). The sampled species included representatives from all continents except Europe (where we had only one sample; Fig. 1) and Antarctica, and from important genera, subgenera, and sections identified in various treatments of morning glories. They include species with localized distributions (e.g., Mexico, Somalia), regional distributions (e.g., South America, Tropical Africa), across regions (e.g., Africa and Asia) as well as pantropical distributions (Supplementary Data 1).

#### i) Herbarium specimens

We obtained permissions to sample mature seeds from herbarium specimens from the Missouri Botanical Garden (MOBOT), Vadense Herbarium (WAG, Netherlands) and the Australian National University Herbarium (ANU), in addition to seeds from personal collections of Convolvulaceae researchers and our own field collections in the United States. Based on the results from the Eich survey^5^, we focused our sampling on species in the tribe Ipomoeeae where all species previously reported to contain EAs occur. Our samples represented a subset of all potential species because most herbarium specimens examined did not have mature seeds. We recorded the most recently annotated name, determined mean seed mass (mg), estimated collection latitude based on location data on the herbarium sheet label and recorded the date of collection. We relied on the species determinations present on the herbarium specimen labels but realize that misidentifications are possible.

#### ii) Field sampling of I. leptophylla and I. pes-caprae

In addition to herbarium specimens, we sampled natural populations of *I. leptophylla* in the northern Great Plains of the United States (Colorado, Nebraska, South Dakota and Wyoming). We identified plants along roadsides in public right-of-ways and collected mature fruits and seeds from 5 – 10 plants per sampling site. Populations of *I. pes-caprae* were sampled in Florida from beaches along both the Atlantic and Gulf coasts. Mature fruits and seeds were collected from extant plants or directly from the sand where dispersed seeds accumulate. For both species, seeds from a single site were combined into a single collection and one seed per population was then randomly selected for EA analysis unless it was less than the 20 mg needed for analysis, where additional seeds were combined until the threshold weight was reached.

### Ergot Alkaloid Analysis

The EA content of seeds was determined by high performance liquid chromatography (HPLC) and liquid chromatography-mass spectrometry (LC-MS). Extracts were prepared by grinding the seeds into a fine powder using a Wiley mill. To prevent cross-contamination, we cleaned the Wiley mill between samples by vacuuming all parts, grinding millet seed (*Pennisetum glaucum*, previously determined to be devoid of EAs), vacuuming again and blowing all parts clean with compressed air. This process was previously tested using known alkaloid-positive samples and alkaloid-negative samples to confirm no detectable cross-contamination. The powdered seed material was then soaked in 1 mL methanol (99.93 % A.C.S. HPLC Grade, Sigma-Aldrich) for three days at 4°C with daily vortexing to promote extraction of EAs. EAs were identified and quantified by methods previously described^50,52^. In short, extracts were separated by reverse-phase HPLC on a C18 column (Prodigy 5-μm ODS3 [150 mm by 4.6 mm]; Phenomenex, Torrance, CA, USA) with a gradient of acetonitrile in aqueous ammonium acetate. Alkaloids were detected with dual fluorescence detectors set at excitation and emission wavelengths of 272 nm/372 nm and 310 nm/410 nm, respectively. Identity of individual peaks was confirmed by LC-MS as previously described^53^.

Based on the results of EA analysis, we classified species as EA+ if EAs were detected in any sample of that species, and as EA− if no EAs were detected in all samples of that species. Some species were included in our phylogeny where we did not have any samples, but were included in Eich^5^. We classified those species as EA+ or EA− following Eich.

### Latitude, Seed Age, Seed Mass, and EAs

We tested for correlation between latitude and EA presence using a logistic regression. Correlation between latitude and EA concentration in EA+ positive species, as well as seed mass, was done using linear regressions. We used the absolute value of each species’ average latitude. EA concentrations and seed mass were averaged for each species. Supplementary Fig. 9 shows variation in seed mass for species with five or more samples. We tested whether EAs decay over time by regressing age on EA concentrations. Seed age was computed from the year 2021 for individual samples with a recorded year of collection (n=135).

### Estimating Morning Glory Phylogeny with ITS Sequence Data

We downloaded the internal transcribed spacer region (ITS) sequence entries from GenBank available in 2020 for 183 out of 210 species that we sampled for EAs (no ITS sequence data were available for 27 of the sampled species). In addition, we added four *Merremia* species to represent outgroup taxa and 23 other important Ipomoeeae species (e.g., *I. batatas*, sweetpotato) previously reported in Eich^5^ (Supplementary Table 1). For each species, we obtained the complete ITS1 + 5.8S rRNA + ITS2 sequence where possible. In some cases, this was obtained by combining partial sequences from multiple accessions of the same species. Relatively little sequence was missing based on comparing the partial sequences against the complete ITS sequences. In total, we obtained 206 sequences including four outgroup species. There were 174 complete ITS1 + 5.8S + ITS2 sequences, 10 sequences from combining partial sequences and 21 were partial ITS1 +5.8S + ITS2 (Supplementary Data 2). The sequence for *I. indivisa* was made available to us from an unpublished dataset (P. Muñoz-Rodríguez). These sequences were aligned using MAFFT v7.450^54^ using the L-INS-i alignment strategy. A maximum-likelihood phylogeny was then estimated using RAxML^55^ implemented in R package “ips” v0.0.11^56^ using the GTR+G+I model, rapid bootstrap analysis, random starting tree and 1000 replicates.

We used a combination of floras and treatments^25–27,57^, biogeography, and results from molecular systematics studies^21,58,59^ to identify 35 evolutionary lineages within Ipomoeeae (comprehensive list of references available upon request). This is not a formal classification but rather our best estimate of identifying clades that represent distinct groups of morning glories on independent evolutionary trajectories. Many of the evolutionary lineages were identified with high confidence given that they corresponded to traditionally identified taxa that could be circumscribed by diagnostic morphological characters and that received strong support as monophyletic groups from molecular systematic studies (e.g., Arborescentes, Batatas, Calonyction, Pharbitis, Poliothamnus)^21,27^. We also identified distinct lineages based on a combination of previous placements in traditional taxa and results from molecular systematic studies (American clades 1, 2, 3; Australian clade; Orthipomoea 1; Eriospermum; Erpipomoea 1, 2; Jalapae). These taxa deserve additional study. Finally, some clades identified here are based on the ITS phylogeny as well-supported monophyletic groups but unite morphologically diverse species from distinct traditional taxa (African clade 5, American clade 4).

We recognize the limitations of identifying distinct evolutionary lineages dependent on a single-gene tree, but 75% of our sampled species had no other gene sequences available. Our results are in general agreement with published multigene phylogenies of a smaller number of species, with some minor differences. For example, in our ITS phylogeny *I. nil* and *I. hederacea* are not sister species, although other evidence suggests that they are sister taxa and perhaps part of a species complex^60^. Similarly, two clades are resolved here that include the Leptophylla group, whereas in other studies they are united in a single clade consistent with their formal treatment based on morphology^61^.

### Testing Phylogenetic Signal for Ergot Alkaloids and Seed Mass

#### i) Ergot Alkaloid Presence

We performed a test of phylogenetic signal to determine whether the distribution of EA+ species across the Ipomoeeae was random or clumped where closely-related species share the same EA status. We used the function “fitDiscrete” from the R package “geiger”^62^ that calculates the likelihood of observing trait values under a model^63^. We selected the ARD (all-rates-different) model as indicated by the lowest Akaike Information Criterion (AICc) score. We then used Pagel’s λ^64^ to assess the phylogenetic signal, which is a robust measure insensitive to taxon sample size and commonly used, facilitating comparisons with other investigations^65^. If λ is equal to or near 1, there is a strong phylogenetic signal. We created a null phylogeny by transforming our phylogeny so that there was no phylogenetic signal (λ = 0) and then compared the likelihood scores and computed the p-value of the two phylogenies to determine if our observed phylogeny significantly differed from the no-signal phylogeny. Additionally, we computed Fritz and Purvis’s D^66^ (FPD) by using the function “phylo.d” from R package “caper”^67^. *D*-statistic values of a trait can be less than 1 (non-random distribution) or greater than 1 (random distribution).

#### ii) Seed Mass

The process for testing phylogenetic signal in seed mass was the same as for EAs except average seed weight for each species was first log10-transformed to normalize the data and the “fitContinuous” function from the R package “geiger”^62^ was used. The FPD statistic can only be applied to binary traits so we did not apply it to seed mass.

### Ancestral Character-State Reconstruction

To assess the distribution and origins of EAs, ancestral character states were estimated using maximum likelihood methods for discrete characters^64^. Specifically, we reconstructed ancestral states for alkaloid presence (yes, no) and seed mass (large, small) using the R packages “ape”^68,69^ and “phytools”^70^. We used “ace” from R package “ape” to fit the best model of trait evolution for our data by comparing AICc scores which shows that ARD (all-rates-different) model was better than other models (equal-rates and symmetric). For each internal node the likelihood that the common ancestor was either EA+ or EA-, or whether the species had large or small seeds, was estimated using a joint estimation procedure^68^. Ancestors were designated as being one state or another if the likelihood of it being that state was greater than 75%.

We also carried out Bayesian stochastic character mapping for EAs^71,72^ via the function “make.simmap” using model “ARD” by running 1000 simulations and summarizing the results using “densityMap”^70^. From this analysis, we estimated for segments of the topology the probability that a particular portion of the lineage was EA+ or EA-. We integrated the results from both analyses to provide our best estimate of the pattern of ancestral character states across the phylogeny, as well as the number of independent origins of the symbiosis.

### Correlated Evolution in Seed Mass and Alkaloid Presence

To test for correlated evolution between seed mass and EAs in morning glory species while accounting for the confounding influence of shared ancestry, we used two methods. First, a phylogenetic regression was performed as implemented in the R package “phylolm”^36^ using log10-transformed seed mass data. Second, Pagel’s test of correlation^35^ was done as implemented in the R package “phytools”^70^ with the function “fitPagel”. Pagel’s correlation assesses correlated evolution between binary characters, so we therefore transformed seed mass into a binary trait. The average seed mass of each species was log10-transformed and the mean of these values was used to separate the large (>1.37, 23.4mg) from the small (<1.37, 23.4mg) group.

### Intraspecific and Interspecific Differences in Ergot Alkaloid Profiles

To evaluate differences in EA profiles between and within species, we used the “phyl.pca” function from the R package “phytools”^70^ to perform phylogenetically-corrected principal component analysis (PCA) for the full dataset of using averaged EA values for each species (Fig. 4a, Supplementary Fig. 7a). We used “vegan”^73^ to perform PCA for the Pes-caprae and ATP clades (Fig. 4b, Supplementary Fig. 7b) using all samples of each species. We did not correct for phylogeny in the clade-specific PCAs as we wanted to evaluate variation between multiple samples of species within the same clade. Due to the chemistry of EAs, only eight out of nine detected EAs were retained for the PCA and further grouped into six distinct EA chemotypes (colored boxes in Supplementary Fig. 1). Ergonovine, lysergic acid α-hydroxyethylamide (LAH), and ergine were combined as lysergic acid amides, while agroclavine was removed from the analysis as it is an intermediate alkaloid product (more details in *Determining Ergot Alkaloid Chemotypes* below). Significance of differences in EA profiles were calculated using analysis of similarity. All EA concentration measurements were transformed into relative concentrations for the analyses.

### Determining Ergot Alkaloid Chemotypes

Table 1 records all EAs quantified in the listed Convolvulaceae-*Periglandula* symbioses. Differences in EAs may be the result of different plant or fungus genotypes but also may result from environmental differences that were not controlled in the diverse herbarium collections sampled. While we present data on the nine EAs detected, we excluded certain EAs in our PCA analysis based on their position and significance in the biosynthetic pathway. Chanoclavine-I is an intermediate product transitional to other EAs except in *I. graminea*, where it was the sole EA detected. The presence of chanoclavine-I alone, indicates genetic differences where the pathway (Supplementary Fig. 1) downstream of chanoclavine-I is halted. Similar logic was applied in excluding agroclavine from PCA analyses. Simple amides of lysergic acid amides were considered as a group, because synthesis of individual lysergic acid amides is not completely independent of one another. While ergonovine can be synthesized independent of other lysergic acid amides^74^, synthesis of LAH is dependent on the genetic capacity to produce ergonovine. Ergine arises as a hydrolysis product from different derivatives of lysergic acid, in particular LAH^75,76^. Its accumulation may be affected by sample age and storage conditions, which were not controlled in the samples analyzed here. Therefore, simple amides of lysergic acid were considered as a single unit when classifying chemotypes. Unique end-products from branches off the clavine portion of the EA pathway, including cycloclavine and lysergol, were considered independent contributors to EA chemotype. Cycloclavine results from a unique combination of alleles in the EA pathway of certain fungi^77^. The biosynthetic origin of lysergol has not yet been determined, but its appearance is sporadic, and it does not serve as an intermediate in the EA pathway^78^. Two ergopeptine alkaloids, ergosine and ergobalansine, were detected in abundance, individually and in combination, in seeds from several EA+ species. Data from *Claviceps purpurea* indicate that different ergopeptine alkaloids result from allelic differences in the gene lpsA^79^. Small quantities of different ergopeptines also may result from permissive binding of alternate amino acid substrates by adenylation domains of peptide synthetases^80,81^.

## Supporting information

All Supplemental Materials

## Statistics and Reproducibility

Statistical analyses used in this study are described in the Methods. Further information on the research design and reproducibility is available via the Nature Research Reporting Summary linked to this article.

## Data Availability

All data necessary to assess the conclusions of this study are available in the main text or the supplementary information. All Supplementary Data can be downloaded from Figshare (https://doi.org/10.6084/m9.figshare.14749512)^82^. All other data are available from the corresponding author on reasonable request.

## Acknowledgements

We thank Eric Knox (Indiana U.), James Solomon and George Yatskievtch (Missouri Botanical Garden), Brendan Lepschi and Kirsten Cowley (Australian National Herbarium) and Jan Wieringa (Vadense Herbarium, Wageningen U.) for help in obtaining seeds from herbarium specimens. We also thank Margaret S. Devall (USDA Center for Bottomlands Hardwood Research) and Kathleen Keeler (U. Nebraska-Lincoln) for seeds from their own collections. We also thank Richard E. Miller (Flower Diversity Institute, Arvada, CO) for his input on interpreting morning glory phylogeny and his recommendations on our phylogenetic analyses. Indiana U. students Michelle McKee, Eric Kimmel, Heather Smith and Clayton Tincher provided lab assistance. W.T.B. was supported by an Anne S. Chatham Fellowship in Medicinal Botany from the Garden Club of America, a grant-in-aid of research from Sigma Xi, an Ecological Society of Australia Student Research Grant and a travel grant from the American Philosophical Society Lewis and Clark Fund for Research and Field Exploration. Additional funding was provided by grant 2012-67013-19384 from the United States Department of Agriculture, National Institute of Food and Agriculture to D.G.P. and grant 429440 from the Simons Foundation to the Smithsonian Tropical Research Institute to K.C. (W. Wcislo, P.I.).

## Contributions

This paper represents part of W.T.B.’s PhD dissertation (2014) in the Department of Biology, Indiana U. W.T.B. and K.C. conceptualized the study. W.T.B., K.C., D.G.P., K.L.R. collected and generated data. Q.N.Q. performed statistical analyses with input from K.C. K.C. led the writing and Q.N.Q. and D.G.P. contributed to writing, reviewing, and editing.

