## Supplementary material for "Diversification of ergot alkaloids and heritable fungal symbionts in morning glories": All Supplemental Materials

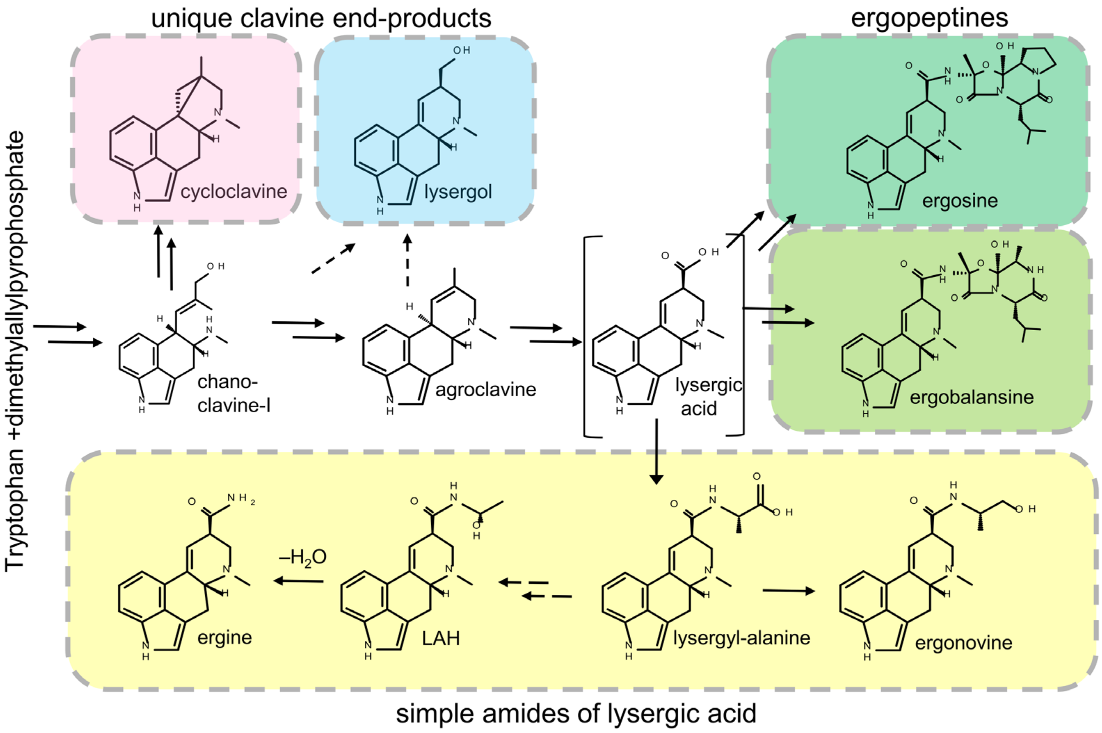


**Supplementary Figure 1.** Diversification of EAs produced in Convolvulaceae-*Periglandula* symbioses. Double arrows indicate one or more omitted intermediates. Dashed arrows indicate uncharacterized steps. Lysergic acid is bracketed to indicate that it is not typically considered a clavine and, as a transient intermediate, typically is not detected in analyses. Colored boxes represent the six distinct EA chemotypes used in PCAs (See Materials and Methods).

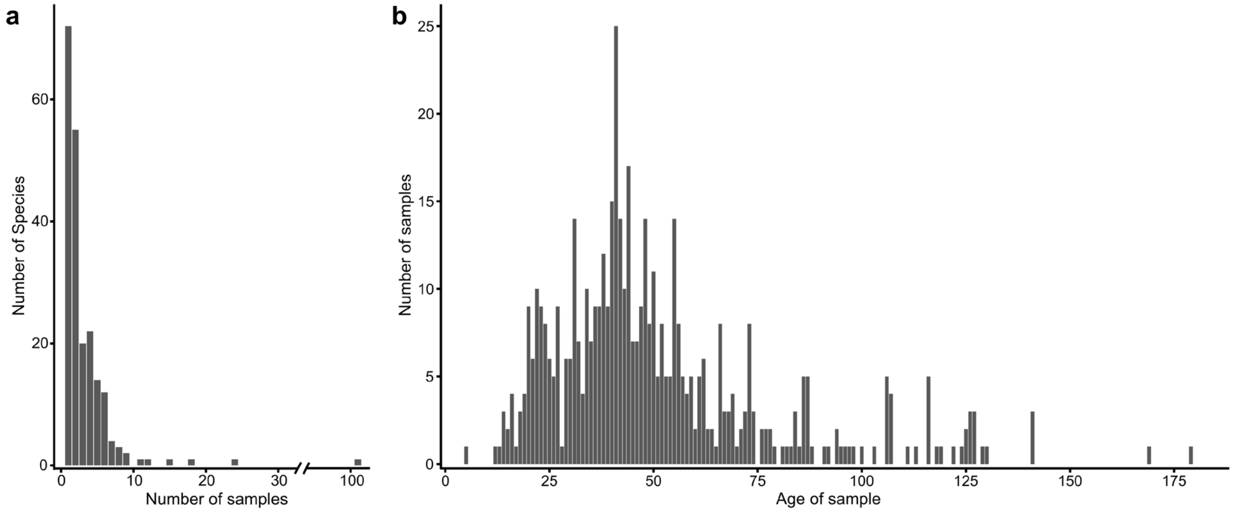


**Supplementary Figure 2. Sample and sample age distribution. a)** Sample distribution among surveyed species; **b)** Sample age distribution among samples with an available year of collection.

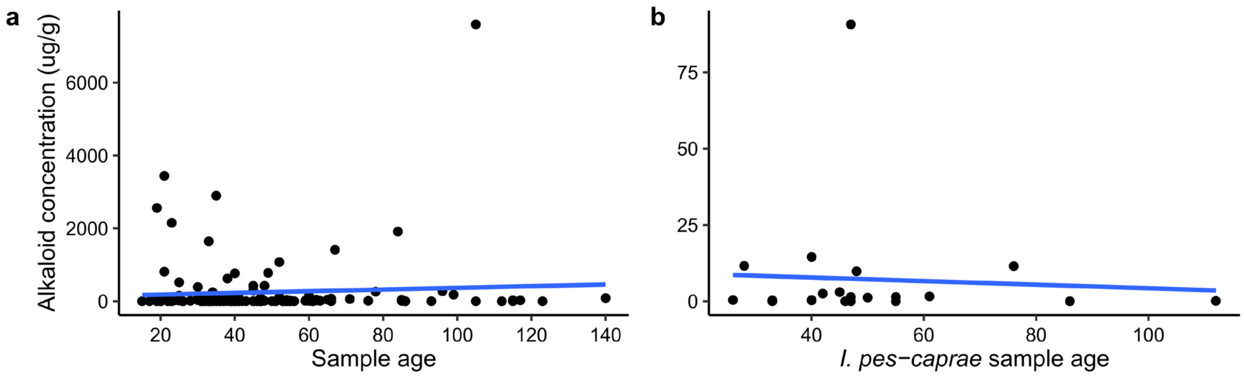


**Supplementary Figure 3.** Ergot alkaloid concentration variation based on sample age in **a)** all EA+ samples, and **b)** *Ipomoea pes-caprae*.

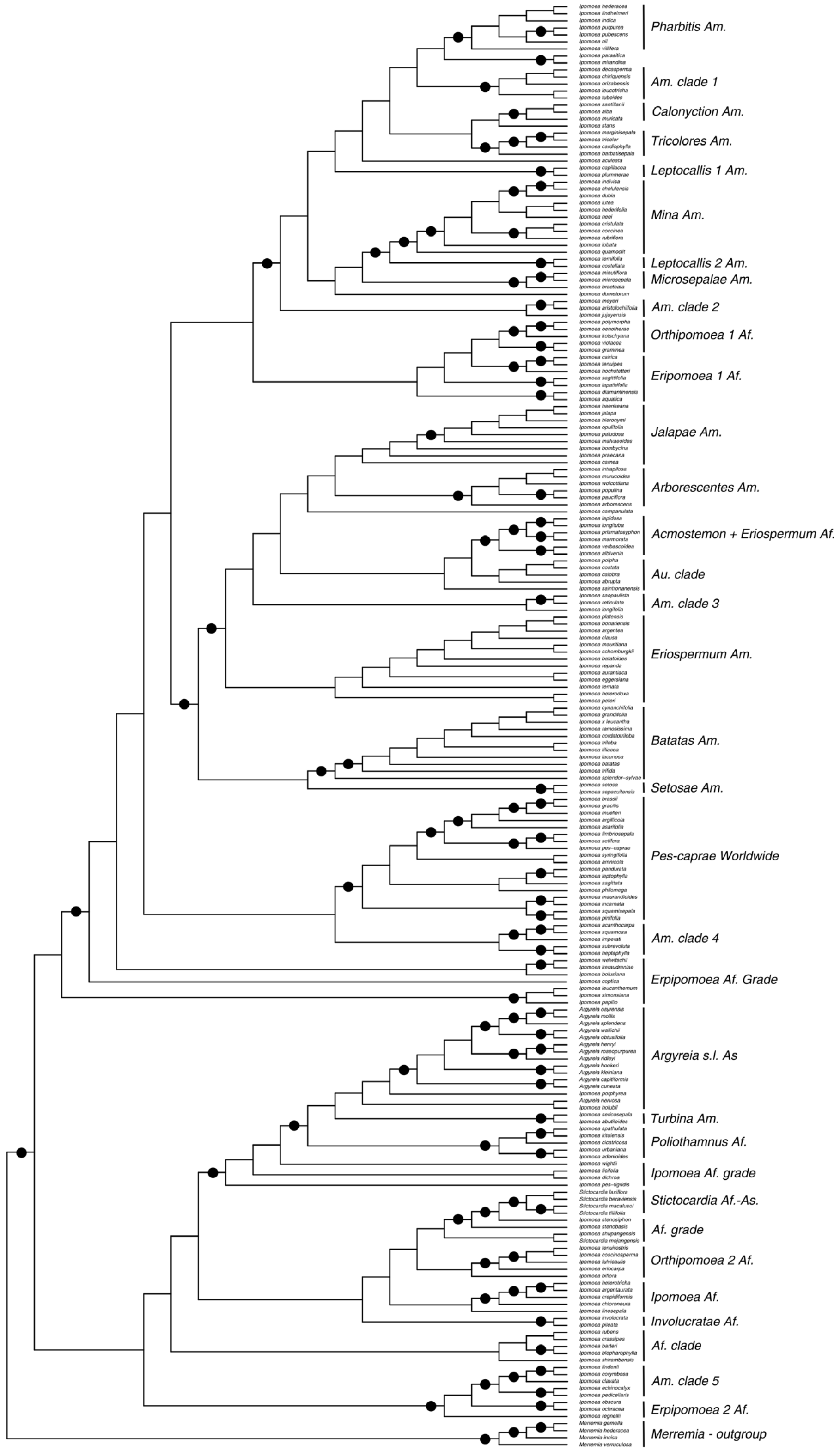


**Supplementary Figure 4.** Maximum likelihood ITS phylogeny with clade denotation. Nodes with bootstrap support >75 are represented by a black circle. Af, African; Am, American; As, Asian; Au, Australian.

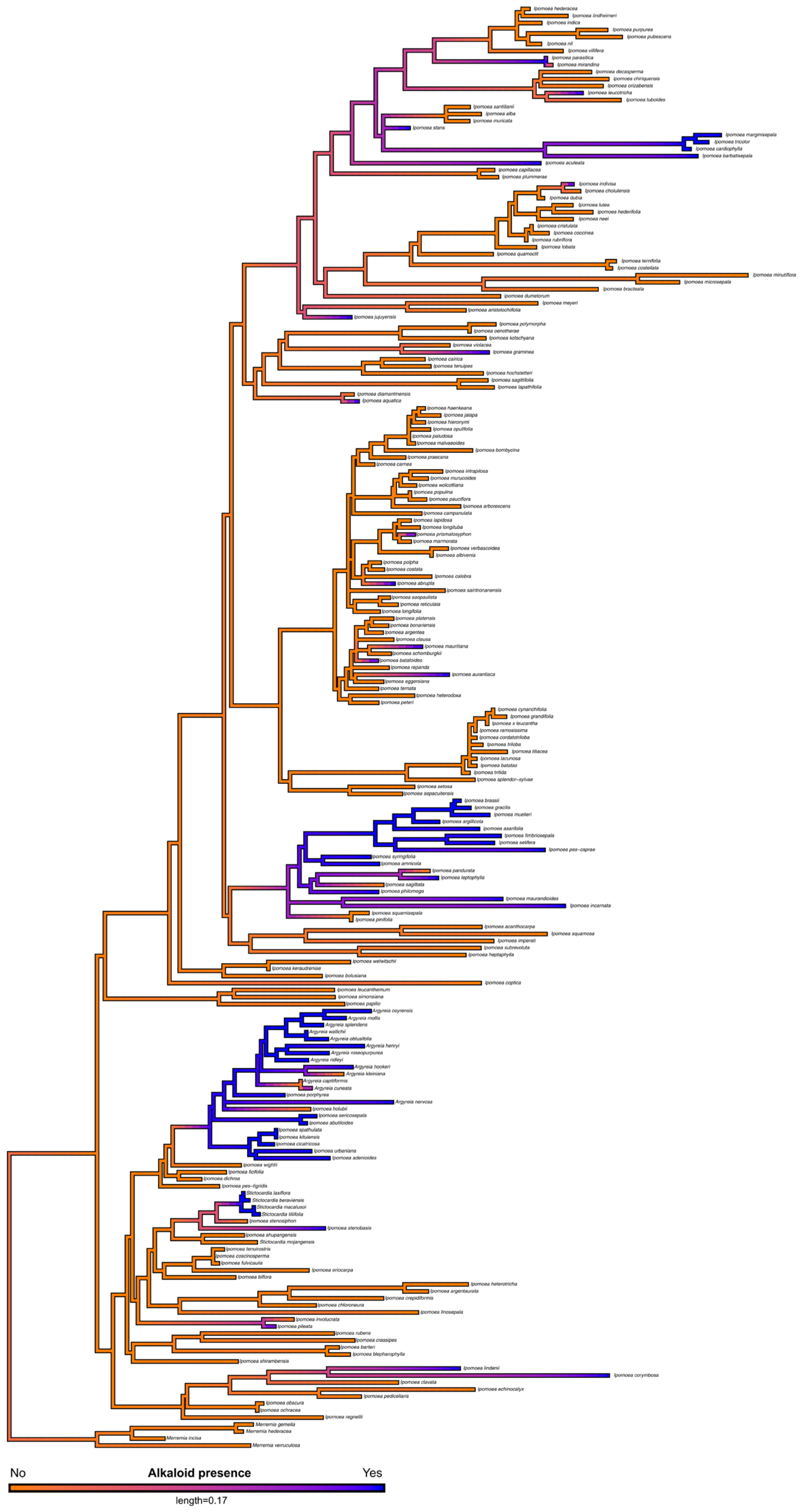


**Supplementary Figure 5.** Density map of Bayesian stochastic character probabilities of alkaloid positive (blue) and alkaloid negative (orange) character states. Legend length equals units of substitution per site.

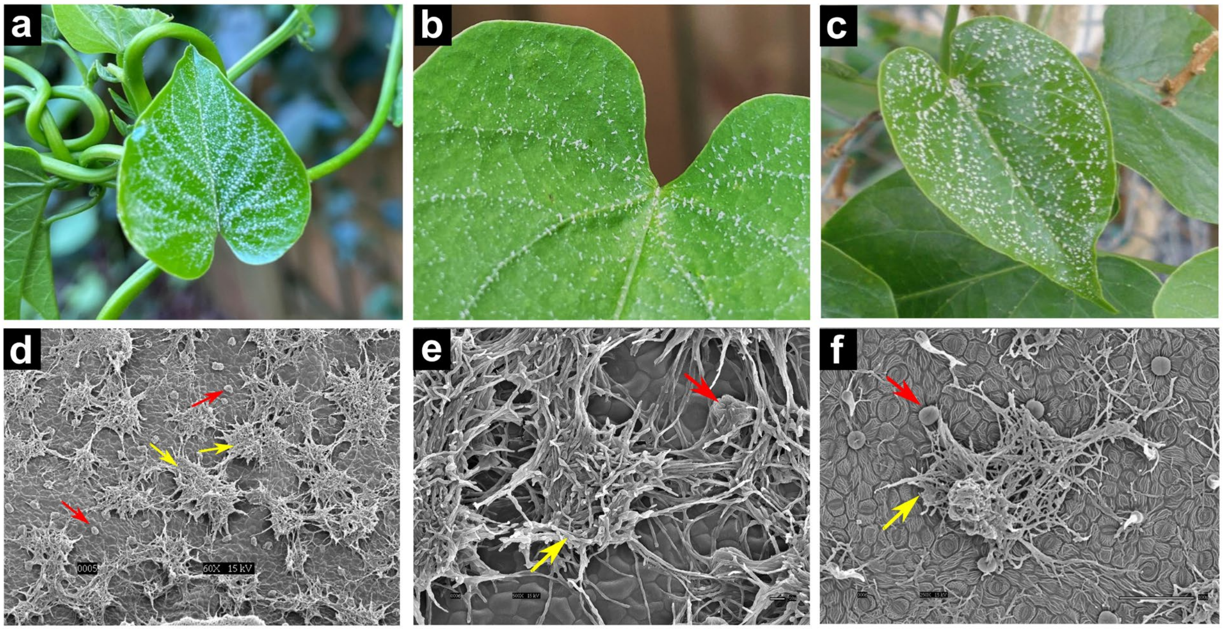


**Supplementary Figure 6. Epiphytic fungal colonies. a,b,c)** Colonies growing along the leaf veins of *Ipomoea corymbosa*. **d,e,f)** Scanning electron microscope images of fungal colonies (indicated by yellow arrows) closely associated with oil glands (indicated by red arrows).

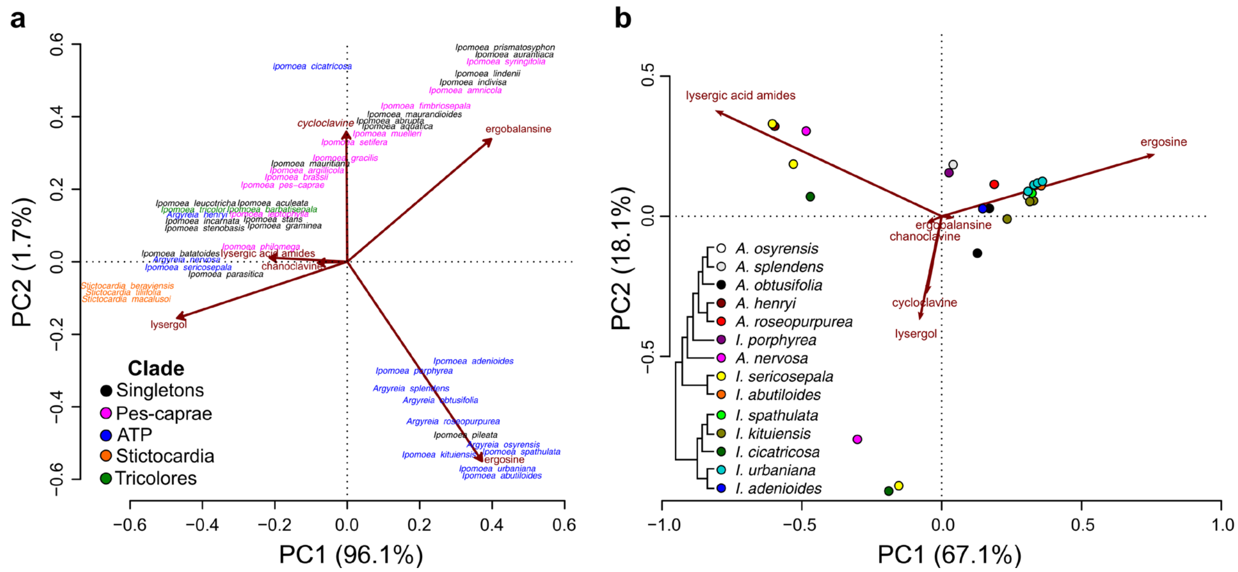


**Supplementary Figure 7.** Clustering of alkaloid chemotypes in alkaloid-positive species. **a)** Phylogenetic PCA showing differences in EA profile of different clades of morning glory, represented by different colors. Alkaloid values are averaged for each species. "Singletons" are clades with only one positive taxa each. **b)** PCA showing differences in EA profiles of different species in the ATP clade.

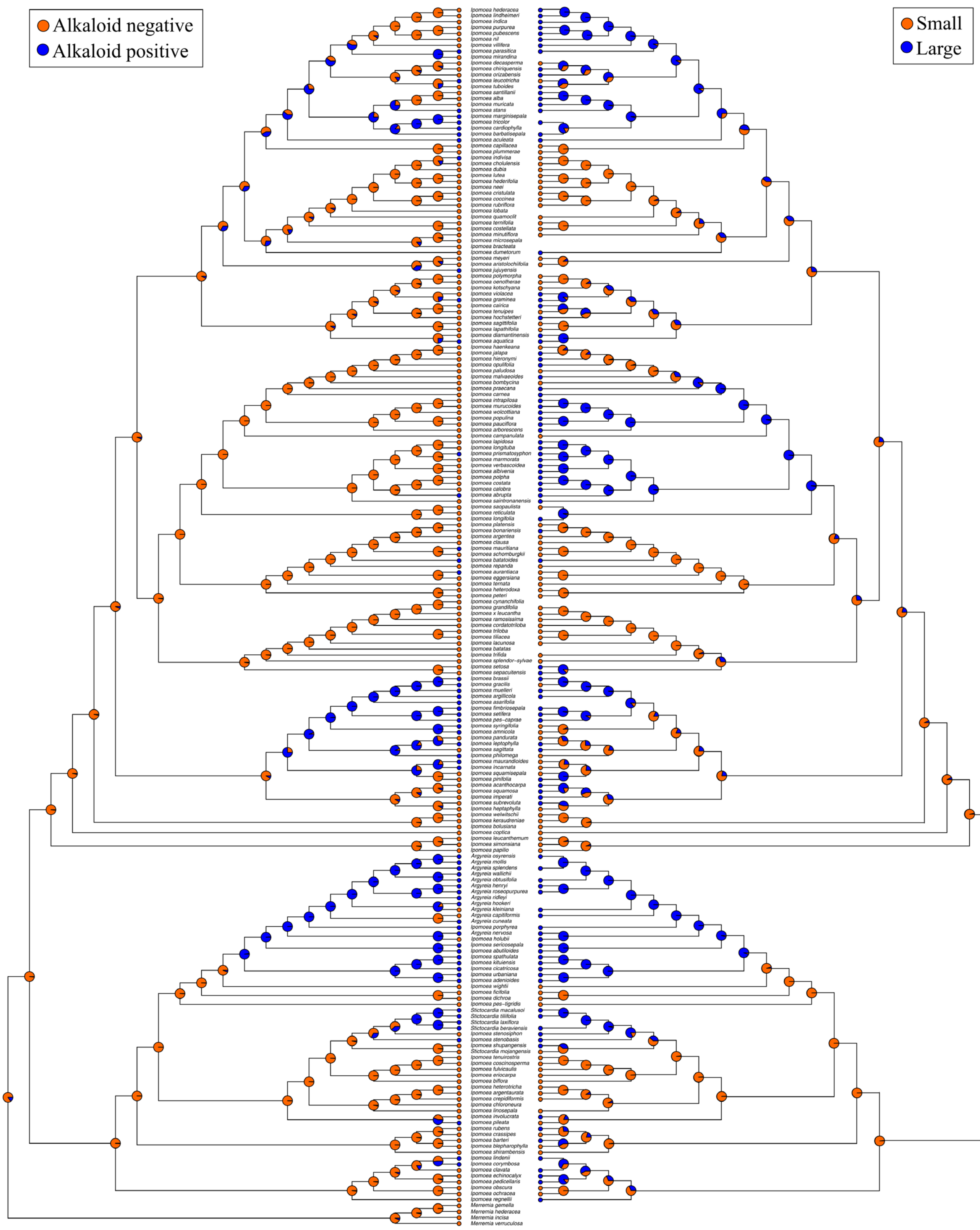


**Supplementary Figure 8.** Comparison of the ancestral state reconstruction of alkaloid presence (left) and seed size (right). Circles at the tips represent current state and pie charts at nodes indicate the relative probability of the ancestral character state.

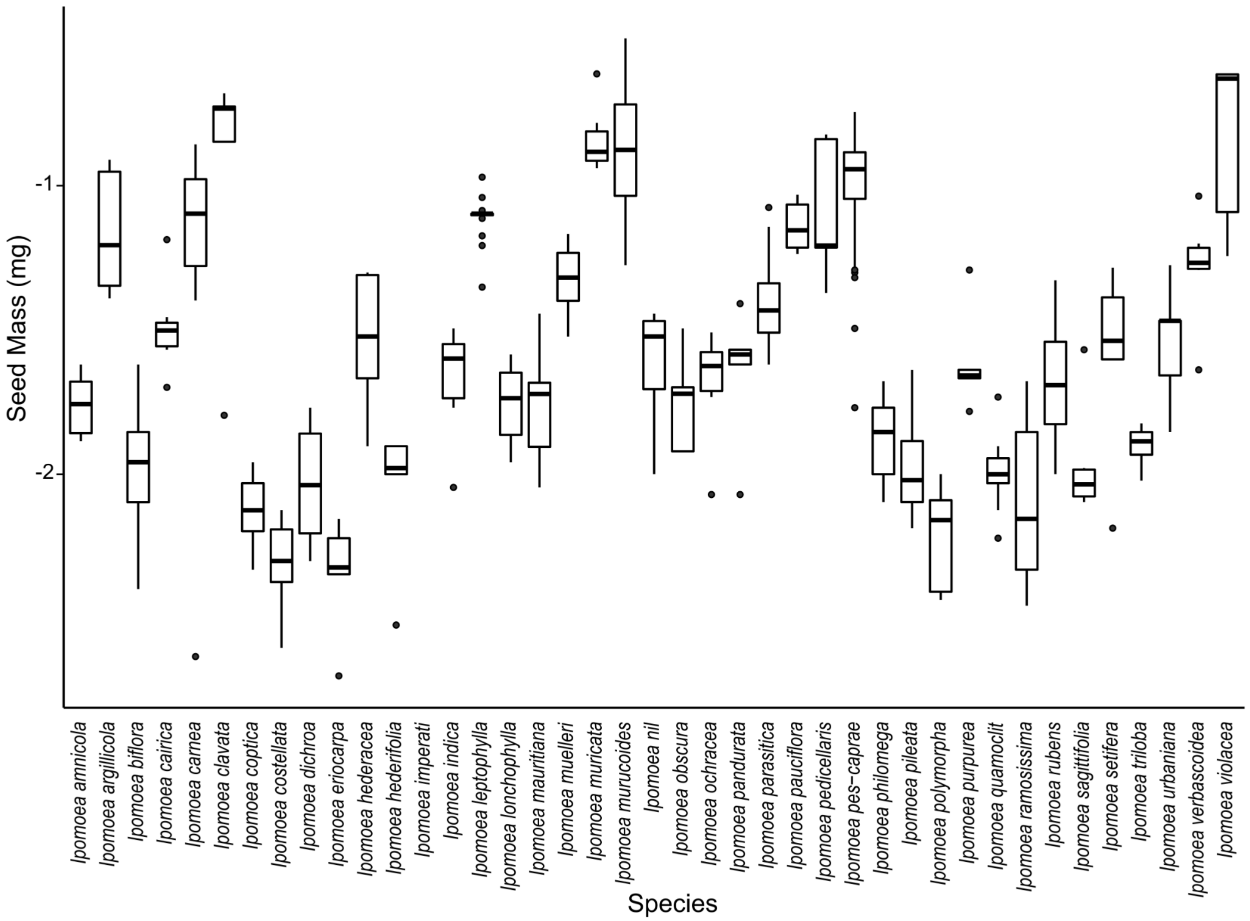


**Supplementary Figure 9.** Seed mass variation in species with five or more samples. Seed mass were log10 transformed.

**Table S1.** Species in Eich (2008) absent from our sampling but are included in the ITS phylogeny.

| Species | Authority | Previous Designation (Eich 2008) | Geographical Distribution |
|---|---|---|---|
| *A. cuneata* | (Willd.) Ker Gawl. | Positive | India |
| *A. hookeri* | C.B.Clarke | Positive | Nepal to Thailand, Andaman Is. |
| *A. mollis* | (Burm f.) Choisy | Positive | Bangladesh to Hainan and Lesser Sunda Is. |
| *A. ridleyi* | (Prain) Ooststr. | Positive | Pen. Malaysia to Sumatera |
| *A. wallichii* | Choisy | Positive | Sikkim to SC. China and N. Indo-China |
| *I. asarifolia* | (Desr.) Roem. & Schult. | Positive | Tropics |
| *I. batatas* | (L.) Lam. | Devoid | |
| *I. bracteata* | Cav. | Devoid | |
| *I. cardiophylla* | A.Gray | Positive | Arizona to Texas and Mexico |
| *I. chloroneura* | Hallier f. | Devoid | |
| *I. corymbosa* | (L.) Roth | Positive | Mexico to Trop. America |
| *I. cynanchifolia* | Meisn. | Devoid | |
| *I. jujuyensis* | O'Donell | Positive | Ecuador to NW. Argentina |
| *I. lobata* | (Cerv.) Thell. | Contradictory | |
| *I. marginisepala* | O'Donell | Positive | S. Bolivia to NW. Argentina |
| *I. microsepala* | Benth. | Devoid | |
| *I. mirandina* | (Pittier) O'Donell | Devoid | |
| *I. reticulata* | O'Donell | Devoid | |
| *S. laxiflora* | (Baker) Hallier f. | Positive | Tanzania to E. South Africa, Madagascar |
